# Children become accurate years before they become fast: electroencephalography dates the separate maturation of decision components

**DOI:** 10.64898/2026.11.17.767330

**Authors:** Han Cui, Yuge Zhu, Bing Li, Shaojia Huang, Tao Xu

## Abstract

Perceptual decisions, a hallmark of cognitive development, become faster and more accurate from early childhood to adulthood, but whether the underlying components mature together, as a single global gain in processing speed would predict, or on separate timetables remains unclear. Behaviour alone cannot distinguish whether age-related improvement reflects clearer sensory encoding, more efficient evidence accumulation, fewer attention lapses or shorter non-decision processes, and prior direct measurements have typically come from small samples spanning limited age ranges. We analysed electroencephalography from 1,534 participants aged 5 to 21 years in the Healthy Brain Network during a contrast-change discrimination task. On the same trials, we measured the visual response and an evidence-accumulation signal, while a decision model estimated accumulation efficiency, attention lapses and non-decision time. We found that the visual response did not grow with age and retained 77% of its strength on missed targets. Instead, accuracy improved through fewer lapses and more efficient accumulation, both reaching half of their developmental change by about 9 years, whereas speed improved later: non-decision time shortened between 11 and 14 years and was longer in children whose accumulation signal rose later. These findings show that perceptual decision-making matures through dissociable stages rather than a single global increase in processing speed: mechanisms supporting accuracy mature in middle childhood, whereas those supporting the fastest responses continue to develop into early adolescence. This framework links developmental theories of processing speed to neural mechanisms of decision-making and provides a reference for identifying which component is disrupted in children who struggle.

**Research Highlights:**

- In 1,534 participants aged 5 to 21 years, children became accurate years before they became fast: accuracy matured 4.6 years before the fastest responses.
- Developmental gains arose after early vision: the visual response did not strengthen with age, and missed targets were registered but not used.
- Decision components follow separate timetables: lapses and evidence accumulation were half mature by about 9 years, non-decision time only by 12.
- These findings challenge the view of processing speed as a single developing capacity and give each decision component a developmental reference for children who struggle.

## Introduction

Children make faster and more accurate decisions about what they see as they grow older, and this improvement is among the most consistent findings in cognitive development. Response times shorten steeply through childhood and more gradually into adolescence, approaching adult levels in the mid-teens (Kail, 1991; Luna et al., 2004), while accuracy keeps rising between 10 and 18 years (Cromer et al., 2015). How the two relate depends on the task: when a response must come before the stimulus ends, slow accumulation of evidence turns into missed responses and lower accuracy rather than into longer response times. Faster processing has been linked to the growth of working memory, reasoning and reading (Ferrer et al., 2013; Fry & Hale, 1996; Kail & Hall, 1994), which is one reason why developmental theories have treated processing speed as a global resource that improves with age (Kail, 1991; Kail & Ferrer, 2007).

Even a simple perceptual choice involves several processes. The brain encodes the stimulus, accumulates the resulting evidence until it suffices to commit to a response, and prepares and executes that response; on some trials the decision process is never engaged. A child can become faster because the stimulus is encoded more sharply, because evidence is accumulated more efficiently, because responses are prepared sooner or because fewer trials are lost to lapses of attention. Which of these processes changes, and when, determines what a developmental gain in processing speed means.

Sequential sampling models separate these contributions formally (O’Connell et al., 2018; Ratcliff & McKoon, 2008). In the diffusion model, the drift rate indexes the efficiency of evidence accumulation (Weigard et al., 2026), the boundary separation indexes response caution, and the non-decision time collects the time spent outside the decision itself, which model estimates do not equate with sensorimotor delay alone (Bompas et al., 2025); a lapse process can be added for trials on which no decision is made (Ratcliff & Tuerlinckx, 2002). Applied to children and adolescents, the model attributes slower responses to differences in several components at once, with a prominent role for lower-quality evidence (Ratcliff et al., 2012), and it has located the processes that differ in children with dyslexia or anxiety (Thompson & Steinbeis, 2021; Zeguers et al., 2011). In 6-to 12-year-olds performing a motion discrimination task, younger children had lower drift rates, wider boundaries and longer non-decision times than older children, and the best model of the children’s data included age effects on drift rate and boundary separation (Manning et al., 2021). In early adolescence, an n-back task in the Adolescent Brain Cognitive Development study showed its largest developmental changes in drift rate and non-decision time (Frogner et al., 2026), and in a stop-signal task faster drift rates went with fewer attention problems whereas non-decision time did not (Wiker et al., 2024).

Electrophysiology adds direct readouts of the stages that these models infer. When stimuli flicker at distinct frequencies, the steady-state visual evoked potential (SSVEP) at each frequency tracks the cortical response to each stimulus, so that the difference between the responses to two competing stimuli indexes the sensory evidence available for a choice (Norcia et al., 2015); such frequency-tagged responses can be recorded reliably in young children (Lochy et al., 2020; Wang et al., 2023). The centro-parietal positivity (CPP) builds up with accumulated evidence, peaks around the response and steepens with stronger evidence (Kelly & O’Connell, 2013; O’Connell et al., 2012; Twomey et al., 2015); its onset and rate can be dissociated (Loughnane et al., 2016), and its build-up rate relates to the speed of perceptual reports (Brosnan et al., 2020). In 6-to 12-year-olds, early occipital and later centro-parietal components both differed from those of adults during motion discrimination (Manning et al., 2019), and the slope of the centro-parietal component related to drift rate (Manning et al., 2021). Preparatory alpha lateralization tracks single-trial response times in 6-to 8-year-olds (Meredith Weiss & Marshall, 2023), and studies of children with dyslexia or attention-deficit/hyperactivity disorder have used SSVEP, CPP and related signals to locate where their decisions differ from those of their peers (Biabani et al., 2025; Stefanac et al., 2021).

Both the visual input to these decisions and the attention that sustains them continue to develop. Younger school-age children detect visual changes less efficiently than older children and adults (Shore et al., 2006), contrast normalization develops through childhood and adolescence (Pei et al., 2017), and the maturation of the alpha rhythm between 5 and 21 years tracks visual white matter and visual detection (Caffarra et al., 2024). The ability to sustain attention improves into adulthood (Fortenbaugh et al., 2015), and between 3.5 and 6 years gains in sustained attention rest more on returning to a task after distraction than on staying on it (Kim et al., 2023).

However, three questions about these gains remain open, and each is hard to answer with the data available so far. The first is where the gain arises. A higher drift rate can come from a sharper sensory representation of the stimulus, or from a more efficient conversion of that representation into a decision variable (O’Connell et al., 2018; Weigard et al., 2026). Behavioural fits cannot tell these two sources apart, because both make responses faster and more accurate. The same question arises in developmental studies of reward, adversity and poverty (Fosco et al., 2022; Vermeent et al., 2024; Zhang et al., 2026). To separate the sources, one needs readouts of encoding and of accumulation on the same trials as the behaviour, yet most developmental tasks do not provide them. Current developmental evidence can thus show that performance improves with age, but not whether the improvement arises before or during the formation of a decision. The second question concerns the targets that children miss. A miss can mean that the stimulus was never encoded, or that it was encoded but the child did not act on the evidence. The answer decides whether lapses belong to the sensory or to the decision side of development. Yet a miss leaves no response to model, and even children’s correct responses can hide processes that never surface in accuracy or response time (Śmigasiewicz et al., 2022). Only a neural readout on the missed trials themselves, as in adults (O’Connell et al., 2009), can therefore determine whether misses in children reflect failed encoding or failed accumulation.

The third question is when each component matures. Here the modelling studies disagree. In 6-to 12-year-olds, the best model needed no age effect on non-decision time (Manning et al., 2021), whereas in early adolescence non-decision time was among the components that changed most (Frogner et al., 2026). Because these studies differ in task, age range and modelling choices, it remains unclear whether the disagreement reflects these differences or different stages of development. The global account also makes a sharper prediction. If one mechanism limits processing speed, the speed of different processes changes along one developmental function (Kail, 1991), so the rate of evidence accumulation and the time spent outside the decision should mature on one schedule, and accuracy and speed with them. This prediction has seldom been tested on the components of the same decisions, although large samples show different trajectories for accuracy and speed (Cromer et al., 2015; Luna et al., 2004). Answering the question is hard for two reasons. First, comparing schedules across studies confounds age with task, so the schedules must be measured within one task across the whole age range. Second, single-child EEG is noisy, and the timing of a signal is harder to estimate than its amplitude (Kiesel et al., 2008; Liesefeld, 2018), so such measurements need large samples. So far, the developmental studies that recorded these signals used modest samples, narrow age ranges or clinical contrasts (Biabani et al., 2025; Manning et al., 2019; Manning et al., 2021; Stefanac et al., 2021). None of them followed sensory encoding, evidence accumulation and model-based decision components together from early childhood to late adolescence within one task.

Here we addressed these questions with the contrast change detection task of the Healthy Brain Network (Alexander et al., 2017; Langer et al., 2017; Shirazi et al., 2024) in 1,534 participants aged 5 to 21 years. In this task, two overlaid gratings flicker at different frequencies while one of them gradually gains contrast, so the SSVEP evidence and the CPP are available on the same trials as the behaviour, within one task across the whole age range (Figure 1). We modelled behaviour with a lapse diffusion model and described each measure by the age at which it completed half of its change between 6 and 16 years, which places behavioural, sensory and decision measures on one developmental time axis. If the gains arise from sharper encoding, SSVEP evidence should increase with age and track drift rate across children; if they arise after encoding, it should stay stable while the decision components change. A missed target that was never encoded should leave little SSVEP evidence, whereas one that was encoded but not accumulated should leave the SSVEP evidence without a CPP. And if one global mechanism limits speed (Kail, 1991), accuracy and speed should mature on a common schedule rather than years apart.

**Figure 1.**
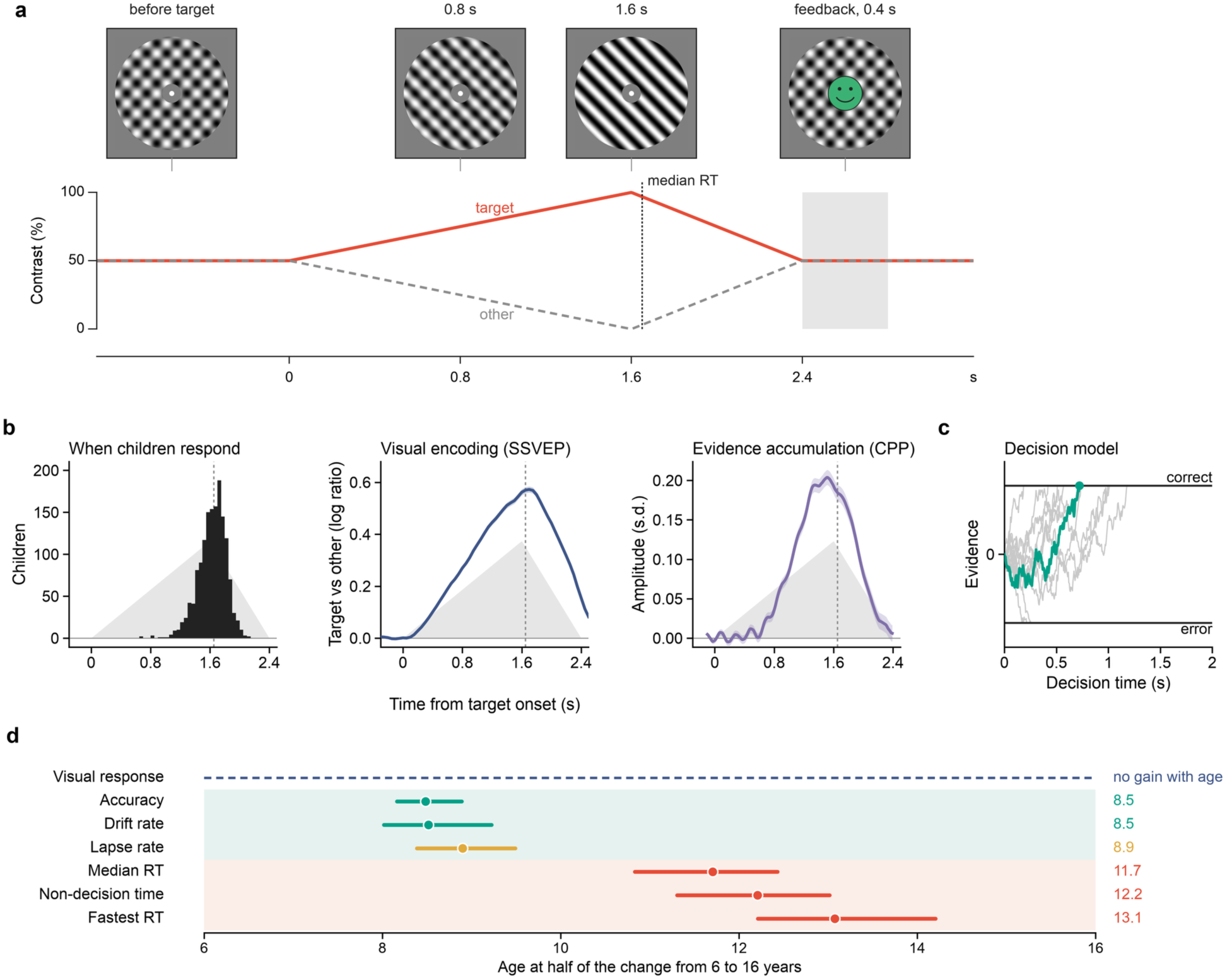
One task, three readouts of the same decision, and their developmental schedules. (a) Contrast of the target grating (left-tilted here) and the other grating during one target, with the screen at four moments; grey bar, feedback. (b) Median RT (n = 1,496), SSVEP evidence (n = 1,505) and CPP (n = 1,504), aligned to target onset; grand averages ± 1 s.e.m. Grey, contrast change; dashed lines, median RT. (c) Diffusion model at the median fitted parameters; green, a correct trial at the median decision time. (d) Age at which each measure completed half of its change from 6 to 16 years, with 95% bootstrap intervals (1,496 children aged 5 to 17 years); the visual response did not increase with age (Figure 3). RT, response time; SSVEP, steady-state visual evoked potential; CPP, centro-parietal positivity; s.e.m., standard error of the mean.

The study makes three contributions. It locates the developmental gain in perceptual decisions after early visual encoding. It shows that the targets children miss are registered by the visual system but not taken up by the accumulation process. And it shows that accuracy and speed rest on decision components that mature years apart, which places earlier modelling studies on one trajectory.

## Methods

### Participants

We analysed the contrast change detection task of the Healthy Brain Network (HBN), a community-referred sample of children and adolescents aged 5 to 21 years from the New York area, recruited through advertisements that encourage participation by families with concerns about psychiatric symptoms in their child. Testing took place at the HBN sites in Manhattan and Staten Island, New York City, USA, an urban setting. The open HBN-EEG release does not include race, ethnicity or socioeconomic status. In a published HBN subsample of 470 youth aged 5 to 17 years, 55% identified as White, 15% as African American, 9% as Hispanic and 17% as multiracial, and household income averaged 3.95 times the federal poverty threshold (SD 2.33; Cohen et al., 2024); these figures describe the population from which our sample was drawn. Of the included participants, 35% were female (Table 1). The recordings came from HBN-EEG releases 1 to 10 on OpenNeuro (Shirazi et al., 2024). Children were included if their age was recorded and they made at least 20 correct responses, the minimum needed to estimate response time (RT) quantiles; 1,534 of 1,666 children met this criterion (Table 1), with exclusions concentrated among 5-to 7-year-olds (19%). EEG measures also required at least 20 valid trials, which 1,505 children provided.

**Table 1.** Sample by age band.

| **Age (years)** | **Recorded** | **Included (excluded)** | **Female (%)** | **EHQ (median)** | **Targets (median)** | **With EEG measures** | **Valid EEG trials (median)** |
|---|---|---|---|---|---|---|---|
| 5–7 | 424 | 342 (19.3%) | 30 | 73 | 72 | 329 | 43 |
| 8–10 | 618 | 589 (4.7%) | 38 | 80 | 72 | 579 | 54 |
| 11–13 | 360 | 348 (3.3%) | 30 | 80 | 72 | 346 | 60 |
| 14–17 | 223 | 217 (2.7%) | 39 | 80 | 72 | 213 | 63 |
| 18–21 | 41 | 38 (7.3%) | 53 | 88 | 72 | 38 | 59 |
| All | 1666 | 1534 (7.9%) | 35 | 80 | 72 | 1505 | 55 |

Included: at least 20 correct responses. EHQ, Edinburgh Handedness Inventory laterality score. Valid EEG trials: correct, RT 0.2–2.5 s and free of amplitude artefacts.

The HBN study was approved by the Chesapeake Institutional Review Board. Participants aged 18 years or older gave written informed consent; for younger participants, legal guardians gave written consent and the participants gave written assent (Alexander et al., 2017). The present study analysed only these publicly available, de-identified data and collected no new data, so no further ethical approval was required.

### Task and EEG recording

Participants fixated the centre of an annulus formed by two overlaid gratings tilted 45° to the left and right of vertical, which phase-reversed at 20 and 25 Hz (Figure 1a; Langer et al., 2017; O’Connell et al., 2012). On each target, the contrast of one grating rose linearly from 50% to 100% over 1.6 s while the other fell to 0%, and both returned to 50% over the next 0.8 s; participants pressed the left or right button to report which grating had increased. Each run contained 24 targets separated by 2.8 to 6.0 s, and the median child saw 72 targets. EEG was recorded at 500 Hz with a 128-channel HydroCel Geodesic Sensor Net (Electrical Geodesics Inc., Eugene, OR, USA); outlying channels were interpolated, and the data were re-referenced to the average and transformed to current source density (Perrin et al., 1989). Trials with eye movements or blinks were excluded from comparisons between trials of the same child; preprocessing settings are listed in the Supplementary Methods.

### Analysis overview

The analysis had five modules that turned each child’s recording into developmental schedules. Modules 1 to 3 read out, on the same trials, the behavioural outcome, the sensory evidence for the choice carried by the SSVEP and the accumulation of that evidence indexed by the CPP (Figure 1b). Module 4 estimated lapse rate, drift rate, boundary separation and non-decision time from the response probabilities and RT percentiles (Figure 1c). Module 5 described each measure as a function of age and related the measures to one another among children of the same age. The output was the age at which each measure completed half of its developmental change, and the relations between the measures.

### Module 1: behaviour

This module turns button presses into the quantities the model explains, keeping the targets that received no response. The response to a target was the first press before the next target that the task scored as correct or incorrect, and targets without such a press were missed. Accuracy was the proportion of correct responses among all targets, so that missed targets lowered accuracy, and the RT distribution was summarized by the 10th, 30th, 50th, 70th and 90th percentiles of correct responses between 0.2 and 2.5 s. The fastest responses were indexed by the 10th percentile.

### Module 2: SSVEP evidence

The visual response itself grows or shrinks with age and recording conditions, so the module has to isolate the part of it that favours the correct choice. Its input was the average of five occipital channels, from which short-time Fourier transforms (400-ms window, 50-ms steps) gave the amplitude at each flicker frequency. On trial *k* at time *t*, the SSVEP evidence was

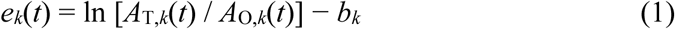

where *A*_T,*k*_(*t*) and *A*_O,*k*_(*t*) are the amplitudes at the frequencies of the target grating, whose contrast increased, and of the other grating, and *b_k_* is the mean of the log ratio before target onset. Positive values mean that the visual response favours the correct choice, and because the measure is a ratio, a change in the overall size of the visual response cancels out (Norcia et al., 2015). The output was each child’s mean SSVEP evidence between 0.8 and 1.2 s after target onset, whose split-half reliability was the Spearman–Brown-corrected correlation between odd and even trials.

### Module 3: centro-parietal positivity

This module reads out how fast and how early evidence is accumulated. Its input was the average of five centro-parietal channels, expressed in units of each child’s trial-to-trial noise so that children with different recording quality were comparable. The single-trial build-up was the slope of the target-locked CPP from 0.2 s after target onset to shortly before the child’s fastest responses, and the response-locked slope was the slope of the child’s average CPP between 0.5 and 0.1 s before the response. The timing of accumulation was the time at which the target-locked average first reached half of its maximum between 0.2 and 1.8 s. Single-child averages are too noisy for this criterion, whereas averages over many children are not, so individual latencies were recovered by jackknifing within age bands (Miller et al., 1998; Smulders, 2010):

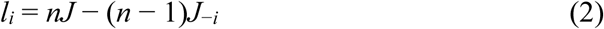

where *n* is the number of children in the band, *J* the half-peak time of the band average and *J*_−*i*_ that of the average without child *i*. The equation converts the small effect of leaving one child out into that child’s own latency. Fractional-area latencies served as a second measure (Kiesel et al., 2008; Liesefeld, 2018).

### Module 4: decision model

A standard diffusion model assumes evidence of constant strength, whereas here the evidence grew with the contrast difference, so we fitted each child’s response probabilities and RT percentiles with a diffusion model whose drift followed the physical stimulus (Ratcliff & McKoon, 2008). Evidence *x* started at zero and evolved as

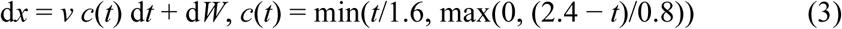

where *v* is the drift rate, *c*(*t*) the contrast difference, rising from 0 to 1 over 1.6 s and returning to 0 by 2.4 s, and d*W* Gaussian noise of unit variance per second. A correct response or an error occurred when *x* reached +*a* or −*a*, provided that the decision time plus the non-decision time *T*_er_ was shorter than 2.5 s. A lapse rate λ captured targets on which no decision was made (Ratcliff & Tuerlinckx, 2002), so that

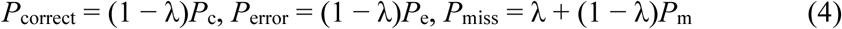

where *P*_c_, *P*_e_ and *P*_m_ are the probabilities of the diffusion process. Because the drift is scaled by the physical contrast, *v* measures how much evidence a child extracts per unit of stimulus change, and λ separates disengagement from slow accumulation as causes of a miss. Parameters were estimated by grid search on simulated predictions, fitting the three response probabilities together with the five RT percentiles. To check that the age effects on *v* and *T*_er_ did not arise from trade-offs between parameters, we refitted the data with *T*_er_ or *a* fixed across children and refitted simulated children whose *T*_er_ followed the observed age curve or the timing of the drift rate; settings, parameter recovery and model predictions are reported in the Supplementary Methods, Supplementary Figures S1 and S2 and Supplementary Note S1.

### Module 5: developmental schedules

Measures in different units have to be compared on one time axis without arbitrary age groups. Age effects were Spearman correlations with age, and each measure was described by a natural cubic spline of age with 4 degrees of freedom, fitted to children aged 5 to 17 years. From the fitted curve *f*, the half-change age was

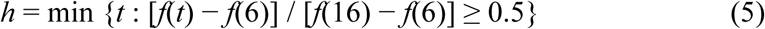

the age at which the measure had covered half of its change between 6 and 16 years. The ratio puts every measure on a common scale from 0 to 1, so that half-change ages are directly comparable. Intervals and differences between half-change ages came from 2,000 bootstrap resamples of children (500 for the refits of simulated children). Relations among children of the same age were partial Spearman correlations controlling age, sex, release and recording quality. We also regressed single-trial log RT on CPP build-up and SSVEP evidence within each child, compared the two signals on correct trials, missed targets and weak-signal correct trials selected in a separate time window, timed the peak of the response-locked CPP, mapped both signals over the scalp, related the four HBN psychopathology scores to the decision components under false discovery rate control (Benjamini & Hochberg, 1995) and compared girls and boys; details are given in the Supplementary Methods.

### Statistical tests

The main tests (Table 2) used one-sided α = 0.05. For the prediction that SSVEP evidence does not increase with age, the upper limit of the 95% interval of its age correlation had to lie below an equivalence bound of ρ = 0.10 (Lakens, 2017). With 1,505 children, a one-sided correlation test had 80% power to detect ρ = 0.06. Age correlations reported without a p value had p < 0.001. Supplementary Table S6 reports the main tests in releases 1 to 4 and 5 to 10 separately. Analyses were run in Python 3.11 with MNE-Python 1.13.2, NumPy 2.4.6, SciPy 1.17.1, pandas 2.2.3 and statsmodels 0.15.0.

**Table 2.** Main tests.

| Test | Prediction | Test statistic | n | Result |
|---|---|---|---|---|
| 1 | Drift rate rises with age | ρ(age, v) > 0, one-sided p < 0.05, with free and with fixed non-decision time | 1,534 | ρ = 0.33; fixed Ter 0.37 |
| 2 | SSVEP evidence does not rise with age and dissociates from drift rate | upper 95% bound of ρ(age, SSVEP) < 0.10, and 95% interval of ρ(age, v) − ρ(age, SSVEP) above 0 | 1,505 | ρ = −0.09 [−0.14, −0.04]; difference [0.34, 0.48] |
| 3 | Missed targets decrease with age | ρ(age, missed-target rate) < 0, one-sided p < 0.05 | 1,534 | ρ = −0.61 |
| 4 | Response-locked CPP slope relates to drift rate at a given age | partial ρ(v, response-locked CPP slope) > 0, one-sided p < 0.05 | 1,505 | partial ρ = 0.09, p < 0.001 |
| 5 | Fast and slow trials differ more in CPP than in SSVEP evidence | within-child Cohen’s d (fast vs slow RT tertile), CPP minus SSVEP > 0, Wilcoxon one-sided p < 0.05 | 1,202 | d = 0.18 vs 0.05, p < 0.001 |
| 6 | Missed targets keep their visual response | lower 95% bound of SSVEP evidence on missed / correct trials > 0.5 | 965 | 0.77 [0.71, 0.83] |
| 7 | Missed targets lack the accumulation signal | CPP on weak-signal correct trials minus CPP on missed targets > 0, Wilcoxon one-sided p < 0.05 | 965 | p < 0.001 |

Partial correlations control a natural spline of log age, sex, release, number of valid trials, noise level and number of interpolated channels. Brackets give 95% bootstrap intervals. Supplementary Table S6 gives the same tests in releases 1–4 and 5–10 separately. ρ, Spearman correlation; v, drift rate; Ter, non-decision time; SSVEP, steady-state visual evoked potential; CPP, centro-parietal positivity; RT, response time; d, Cohen’s d.

### Design choices

Three choices distinguish this analysis from earlier developmental studies. Same-trial readouts assign a change in drift rate to encoding or to the steps that follow it, which behaviour alone cannot do. Scaling the drift by the contrast ramp and adding a lapse rate match the model to gradual targets that children sometimes leave unanswered; a constant drift without lapses would attribute these misses and the late responses on the ramp to slow accumulation. Half-change ages place components with different units on one developmental time axis, which age-group comparisons cannot do.

## Results

### Accuracy improved years before speed

We first asked whether accuracy and speed improve together. Between 6 and 16 years, accuracy rose from 56% to 89% (ρ = 0.56) and missed targets fell from 27% to 5% (Figure 2a; Table 2). Responses also became faster (median correct RT: ρ = −0.38), but the speeding was smaller and came later: the fastest responses, indexed by the 10th percentile of each child’s RT distribution, did not shorten before the age of 11 (Figure 2b). Accuracy completed half of its change between 6 and 16 years at 8.5 years, but the fastest responses only at 13.1 years, a lag of 4.6 years (95% CI [3.6, 5.8]; Figure 2c,d). The same pattern held when missed targets were left out: accuracy among answered targets rose from 79% to 93% and completed half of its change at 8.4 years, 4.7 years before the fastest responses. Median correct RT also lagged accuracy, by 3.2 years (Supplementary Table S4).

**Figure 2.**
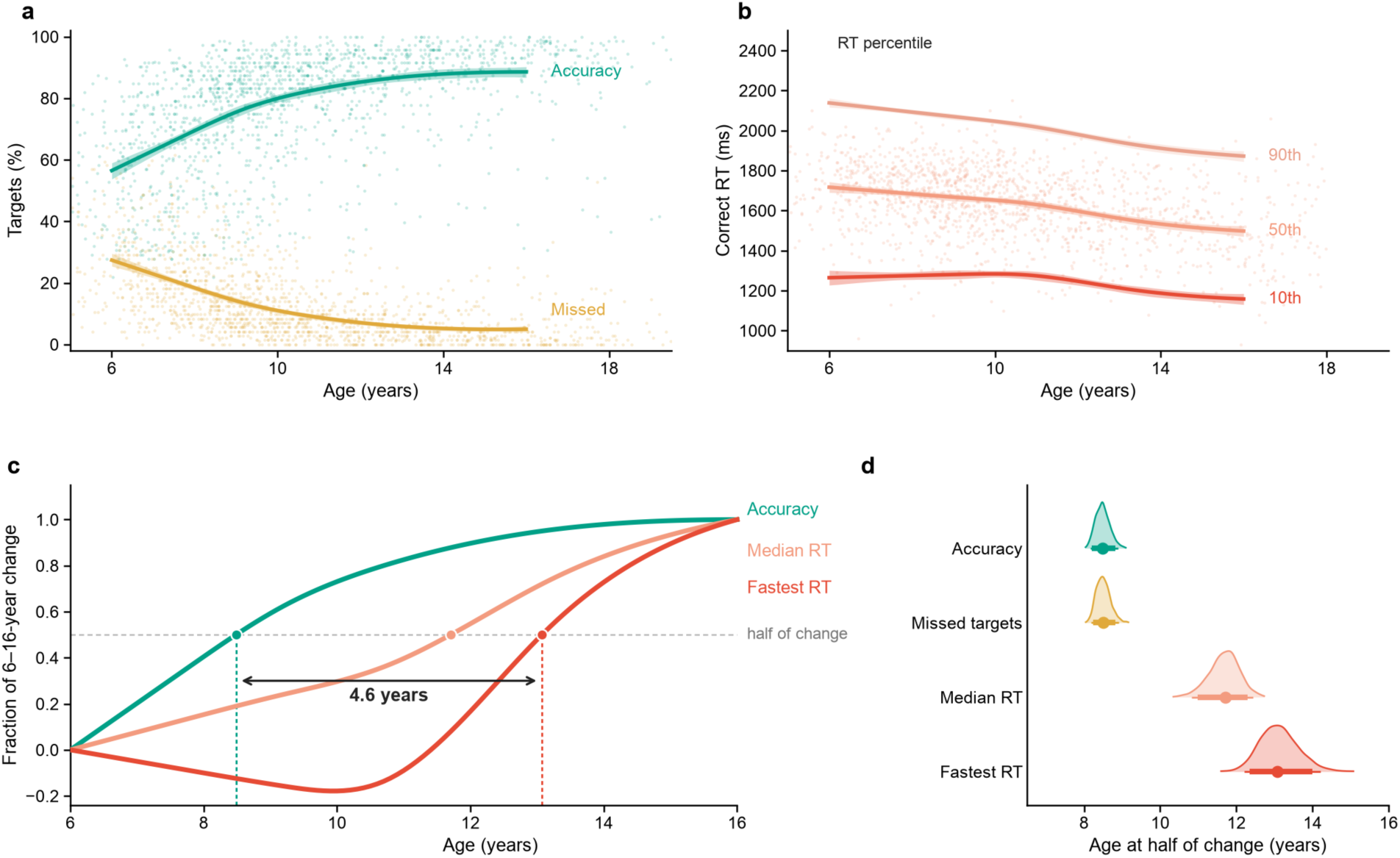
Children became accurate before they became fast. (a) Accuracy (correct responses among all targets) and missed targets by age. Dots show children; lines and bands show natural spline fits and 95% bootstrap intervals. (b) 10th, 50th and 90th percentiles of each child’s correct RT. (c) Each measure expressed as the fraction of its change from 6 to 16 years; circles mark the age at which half of the change was complete. (d) Bootstrap distributions of these half-change ages; thick and thin bars show 90% and 95% intervals. Fits and half-change ages use the 1,496 children aged 5 to 17 years. RT, response time.

### The early visual response did not strengthen with age

To locate the gain, we then examined the early visual response. The SSVEP evidence followed the physical contrast difference in every age band (Figure 3a) but did not increase with age (ρ = −0.09; Table 2). The upper bound of its 95% interval lay below the equivalence bound of 0.10, and the age effect on drift rate exceeded that on SSVEP evidence. Scalp maps showed similar occipital SSVEP evidence in every age band (Supplementary Figure S3). Over the years in which drift rate rose by about one standard deviation, SSVEP evidence fell by 0.40 standard deviations, mainly in mid-adolescence (Figure 3b), so the visual response moved against the gain in decision efficiency. Its split-half reliability was 0.87 to 0.91 in every age band from 5 to 17 years, and its onset did not differ between young children and adolescents (Figure 3c,d). At a given age, weaker SSVEP evidence went with more lapses but not with a lower drift rate (Figure 3e). Differences in decision efficiency, whether between ages or between children of the same age, were thus not explained by early visual encoding.

**Figure 3.**
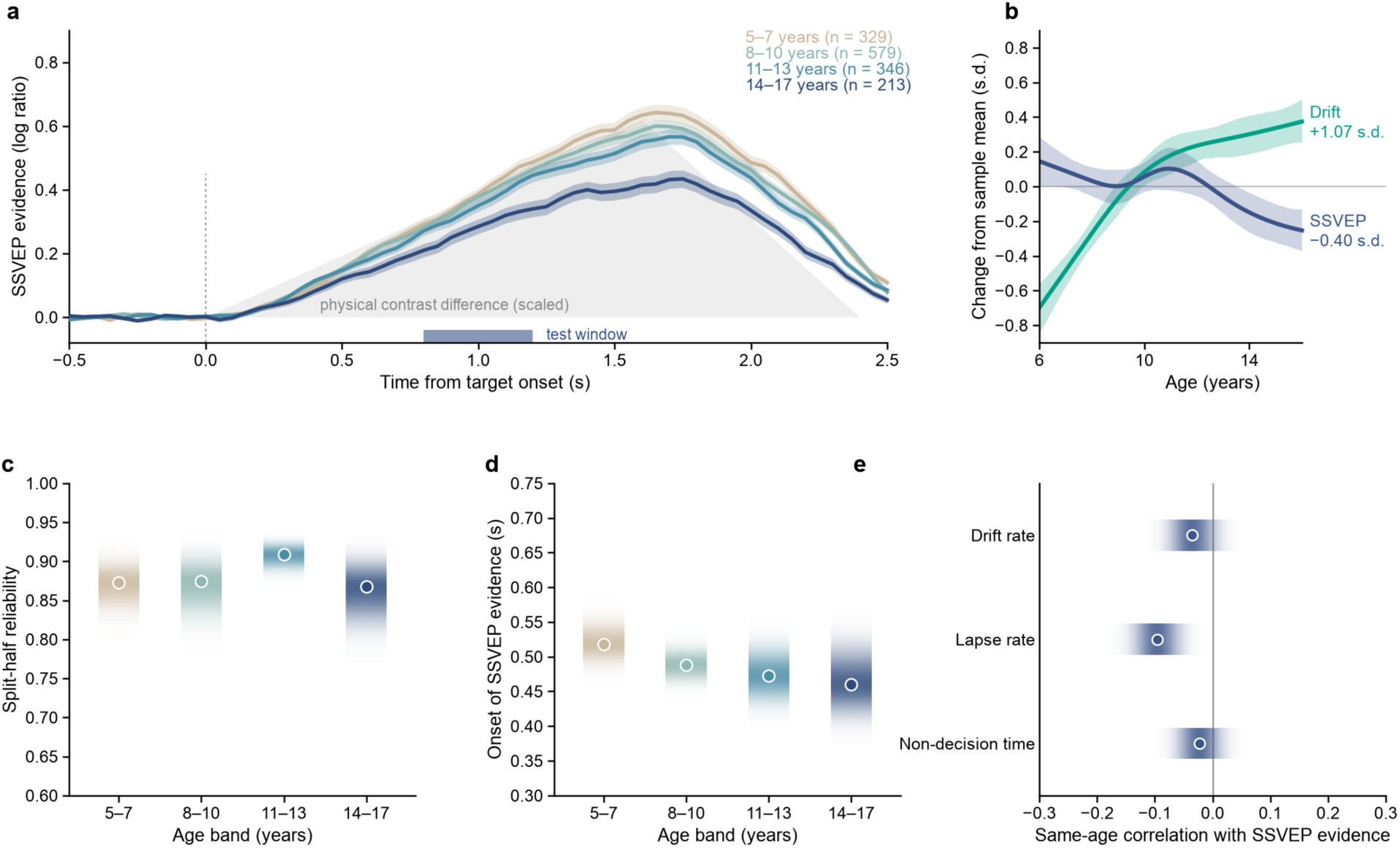
The early visual response to the changing grating did not strengthen with age. (a) SSVEP evidence (log ratio of the responses at the target and non-target flicker frequencies, baseline-corrected) by age band; bands show ±1 s.e.m. The grey area shows the physical contrast difference, scaled; the bar marks the 0.8–1.2 s test window. (b) SSVEP evidence and drift rate by age, both in standard deviations of the sample; bands show 95% bootstrap intervals. (c) Split-half reliability of the SSVEP evidence in each age band. (d) Time at which the band-average SSVEP evidence reached 25% of its maximum. In c and d, shading shows bootstrap distributions. (e) Same-age partial correlations of SSVEP evidence with drift rate, lapse rate and non-decision time, with 95% intervals. SSVEP, steady-state visual evoked potential; s.e.m., standard error of the mean.

### Lapses, drift rate and non-decision time changed with age

To find which decision components carried these gains, we fitted the lapse diffusion model, which attributed them to three components. With age, the lapse rate fell from 19% to 5% and drift rate increased by about 80%. Non-decision time shortened from 940 to 740 ms (Figure 4a–c), while boundary separation increased slightly (Supplementary Table S2). In counterfactual predictions, fewer lapses and higher drift rates together carried the gain in accuracy. Faster responses reflected both a higher drift rate and a shorter non-decision time, which each accounted for about half of the age-related speeding when boundary separation was held constant (Supplementary Note S1). The model reproduced the data in every age band and recovered its parameters and their age effects in simulation (Supplementary Figures S1 and S2).

**Figure 4.**
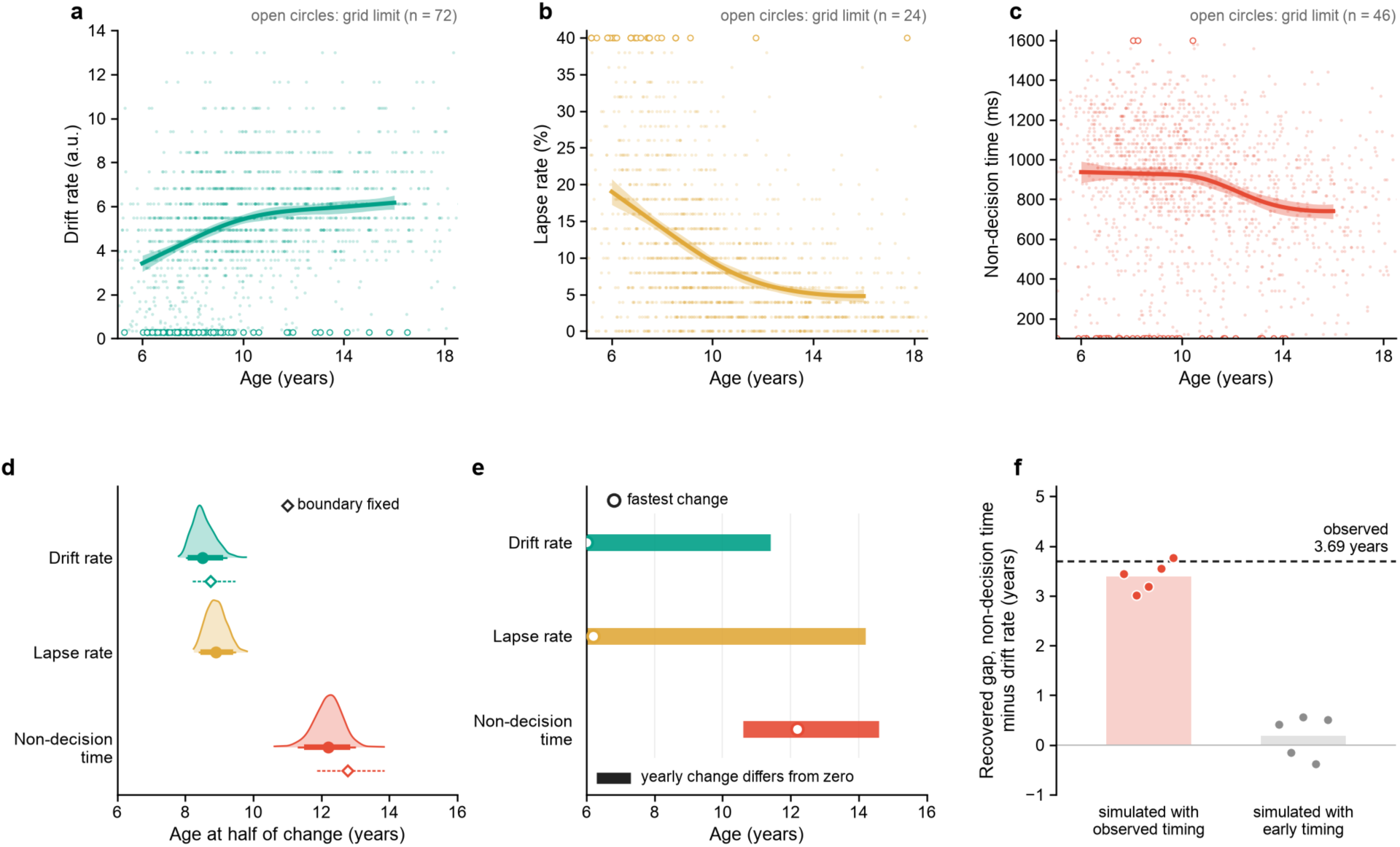
Decision components that follow encoding matured on two schedules. (a–c) Drift rate, lapse rate and non-decision time by age, with spline fits and 95% bootstrap bands. Open circles mark estimates at a limit of the parameter grid. (d) Age at which each component completed half of its change from 6 to 16 years (half-eye; 90% and 95% intervals) and refits with the decision boundary fixed (diamonds). (e) Ages at which the yearly change differed from zero (95% bootstrap); circles mark the fastest change. (f) Gap between the half-change ages of non-decision time and drift rate recovered from simulated children whose non-decision time followed the observed timing or matured with drift rate; dots show five simulation runs, the dashed line the observed gap.

### Within children, RT followed the accumulation signal more closely than the visual response

We then turned to single trials. There, a steeper build-up of the CPP predicted a faster response, whereas SSVEP evidence on the same trials predicted RT about a quarter as strongly (n = 1,202; Figure 5b). The difference between the two weights was −0.047, so RT followed accumulation more closely than the visual input. The CPP–RT coupling did not change with age (ρ = −0.05, p = 0.10; Figure 5e). Between children of the same age, the response-locked CPP slope was weakly related to drift rate (partial ρ = 0.09; Table 2).

**Figure 5.**
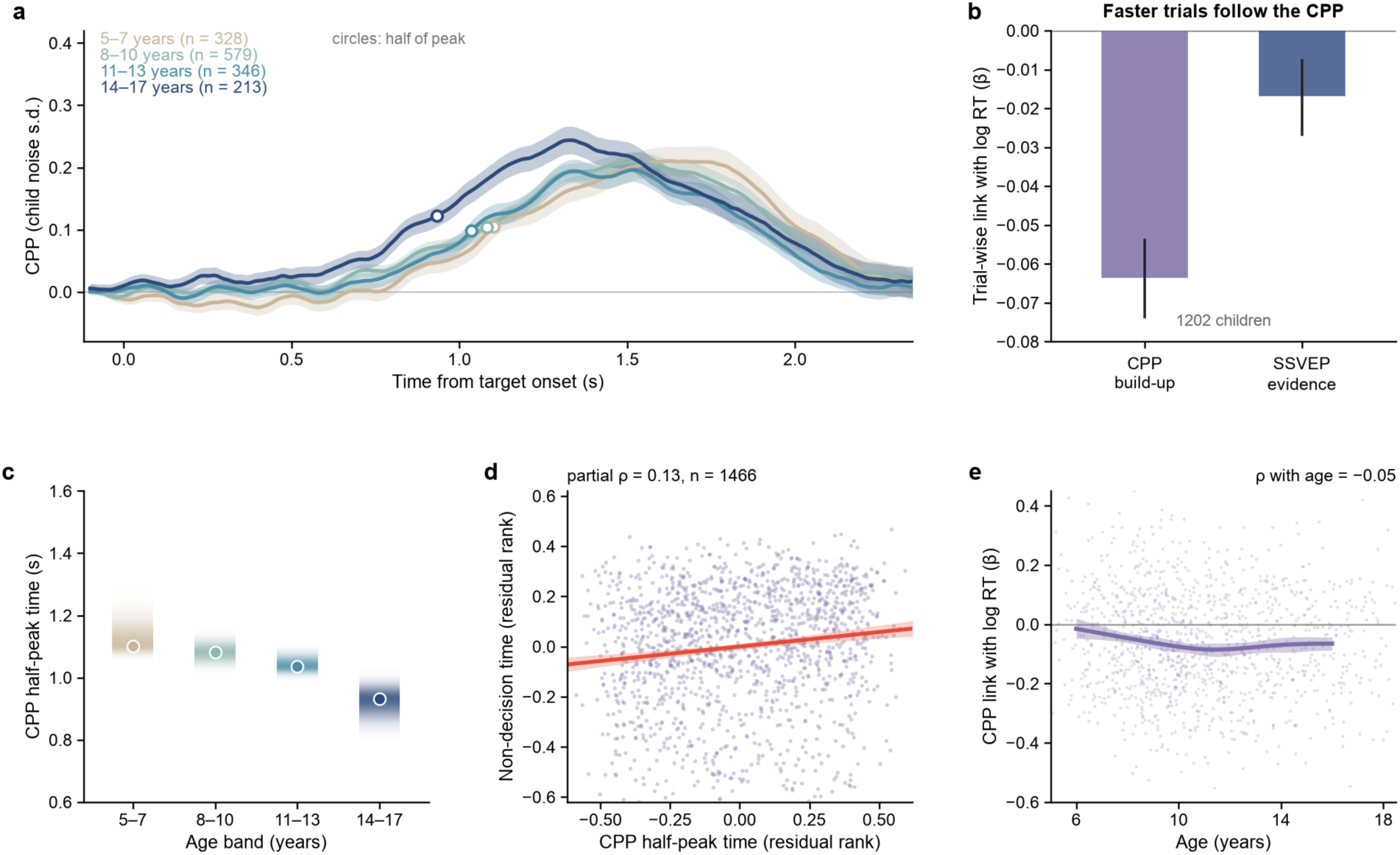
The centro-parietal accumulation signal followed RT within children, and its timing went with non-decision time. (a) Target-locked CPP on correct, eye-movement-free trials by age band, in units of each child’s trial-to-trial noise; bands show ±1 s.e.m., circles mark half of the peak; band sizes count children with valid eye-movement-free CPP trials. (b) Within-child regression weights of log RT on single-trial CPP build-up and SSVEP evidence, averaged over 1,202 children; bars show 95% bootstrap intervals, and the two weights differed by −0.047 (95% CI [−0.060, −0.033]). (c) Half-peak time of the band-average CPP with bootstrap distributions. (d) Same-age relation between jackknifed half-peak time and non-decision time, after controlling age band, overall RT and recording covariates (residual ranks); the line shows the fit with a 95% bootstrap band; children aged 5 to 17 years are shown (n = 1,466). (e) Within-child CPP regression weight by age. CPP, centro-parietal positivity; RT, response time; SSVEP, steady-state visual evoked potential; CI, confidence interval; s.e.m., standard error of the mean.

### The timing of the accumulation signal went with non-decision time

Because speed improved late, we next asked when the accumulation signal rises. The band-average CPP reached half of its peak 1.10 s after target onset at 5 to 7 years and 0.93 s after onset at 14 to 17 years (Figure 5a,c). At a given age, children whose jackknifed half-peak time was later had longer non-decision times, even after age band and overall RT were controlled (partial ρ = 0.13, n = 1,504; Figure 5d; Supplementary Table S3). A later rise of the accumulation signal thus went with a longer non-decision time. The relation held for fractional-area latencies but did not extend to drift rate (Supplementary Table S3). Scalp maps showed a focal central positivity by 0.8–1.0 s at 14 to 17 but not at 5 to 7 years (Supplementary Figure S3). At a given age, non-decision time was unrelated to the interval from the response-locked CPP peak to the press (partial ρ = −0.05; Supplementary Table S3).

### Missed targets were registered visually but not taken up

To identify what a missed target is, we compared missed and correct targets in the 965 children with enough trials free of eye movements. SSVEP evidence on missed targets remained at 77% of its level on correct trials, whereas the CPP stayed close to its pre-target level (Table 2). We selected each child’s weakest third of correct trials by SSVEP evidence at 0.6–0.8 s and measured them in a later window, 1.2–1.4 s. On these trials SSVEP evidence was no higher than on missed targets (difference −0.05, n = 871), yet the CPP was higher (difference 0.06; Figure 6; Supplementary Table S7). Selecting late and measuring early gave the same result (Supplementary Table S7). Scalp maps showed occipital SSVEP evidence on missed targets but no central build-up (Supplementary Figure S3). Missed targets were therefore registered by the visual system but not taken up by the accumulation process.

**Figure 6.**
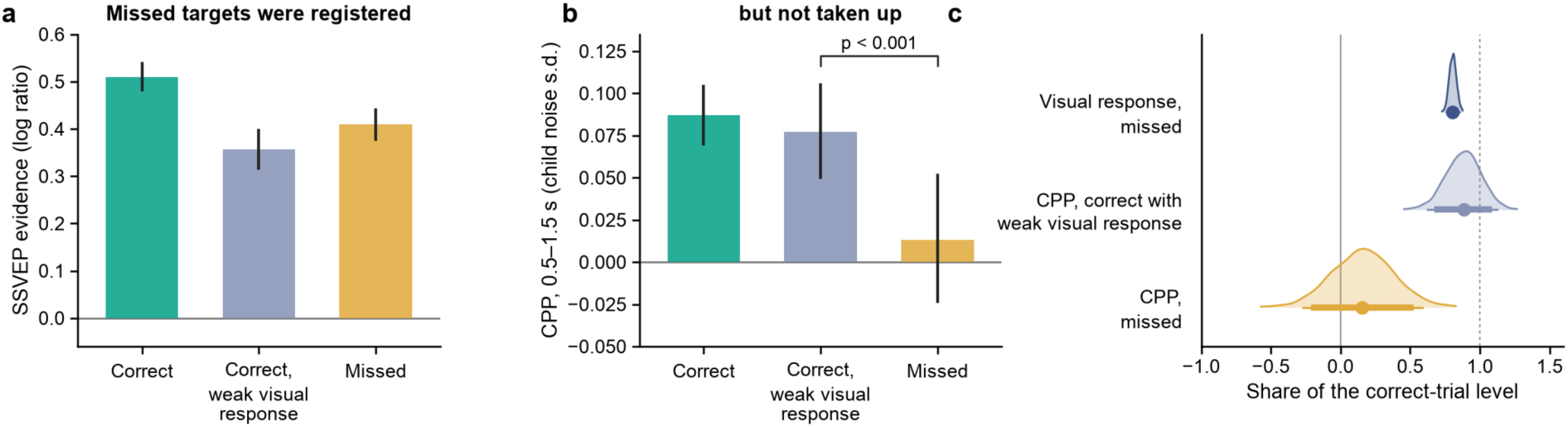
Missed targets were registered but not taken up by the accumulation signal. (a) SSVEP evidence at 1.2–1.4 s and (b) CPP (0.5–1.5 s after target onset) on correct trials, correct trials with a weak visual response (each child’s lowest third of SSVEP evidence at 0.6–0.8 s) and missed targets. Bars show means across children with 95% bootstrap intervals; p, one-sided Wilcoxon signed-rank test. (c) Bootstrap distributions of each signal on missed targets or weak-signal correct trials as a share of its level on correct trials; thick and thin bars show 90% and 95% intervals. n = 871 children with at least 3 missed and 15 correct trials free of eye events and amplitude artefacts in both windows. SSVEP, steady-state visual evoked potential; CPP, centro-parietal positivity.

### Drift rate and lapses matured before non-decision time

To test the global account, we compared the components’ schedules. Drift rate and lapse rate completed half of their change between 6 and 16 years at 8.5 and 8.9 years, whereas non-decision time did so at 12.2 years (Figure 4d). This lag of 3.7 years (95% CI [2.6, 4.7]) shows that the components did not mature together. Drift rate changed fastest before the age of 11, while non-decision time changed little before 10 years and then shortened mainly between 11 and 14 years (Figure 4e). This timing explains why the fastest responses did not shorten before 11 years. A higher drift rate mainly shortens slow responses, and between 5 and 10 years median correct RT fell by about 70 ms while the 10th percentile did not change. The lag persisted with boundary separation fixed across children (4.0 years). When we refitted simulated children whose non-decision time matured together with their drift rate, the recovered lag was only −0.4 to 0.6 years (Figure 4f); alternative curve settings gave similar lags (Supplementary Table S4). These model-based schedules parallel the model-free finding that accuracy improved years before the fastest responses.

At a given age, psychopathology scores were at most weakly related to the decision components (all |partial ρ| ≤ 0.08, none surviving false discovery rate control), and the age effects were unchanged when attention problems and the general psychopathology factor were controlled and in children scoring below the median on both (Supplementary Table S5). Half-change ages did not differ between girls and boys.

## Discussion

Our aim was to identify where and when perceptual decisions become more efficient between early childhood and late adolescence: whether children improve by seeing more clearly or by judging better, whether missed targets go unregistered or unused, and whether the decision components mature together or years apart. To answer them, we read out the visual response, the accumulation signal and the decision components on the same trials of 1,534 participants and dated each measure by the age at which it completed half of its change between 6 and 16 years. The visual response did not strengthen with age, so children improved by judging better. Missed targets point the same way: the visual response remained while the accumulation signal stayed flat, so they had been registered but not used. Nor did the components mature together, as one global gain in speed predicts: accuracy, the lapse rate and the drift rate reached their half-change age at about 9 years, the fastest responses and non-decision time only at 12 to 13 years (Figure 1d). These findings change the account of how decisions develop in three ways. First, they place the lower-quality evidence that diffusion modelling attributes to children (Ratcliff et al., 2012) after the early visual response, in the conversion of that response into a decision variable. Second, they show that a missed target is lost after the visual system has registered it. Third, they split processing speed, often treated as one capacity that changes along a single developmental function (Kail, 1991), into components that mature years apart.

Earlier work located part of children’s immaturity in the visual response: Manning et al. (2019) found an additional positive peak in children’s early occipital response to coherent motion and read it as a difference in stimulus encoding. Here, the visual response did not strengthen with age. Both can hold because Manning et al. described the shape of the response, whereas our SSVEP evidence, a log ratio of the two grating responses, removes age differences in overall response magnitude and isolates the evidence on which the choice depends. The shape can thus change with age while the evidence for the choice does not grow, which moves the developmental difference from the strength of encoding to how the available evidence is converted into a decision.

In adults, momentary lapses begin with reduced prestimulus activity in attentional control regions and are accompanied by weaker sensory responses (Weissman et al., 2006), whereas in a sustained attention task the SSVEP before missed targets can match that before detected ones (O’Connell et al., 2009). Children’s missed targets resembled the second case, failing after encoding but before accumulation. Attentional re-engagement offers a developmental account. Young children are more easily captured by distracting events than older children and adults (Volkmer et al., 2022), and their gains in sustained attention rest more on returning to a task than on staying on it (Kim et al., 2023). What develops, on this view, is how quickly a registered target is taken up for a decision. Training-related reductions in children’s response variability have likewise been traced to evidence accumulation and attributed to attention control (Cubillo et al., 2023).

Why, then, did speed mature years after accuracy? Non-decision time collects every delay outside the decision (Bompas et al., 2025), so its late change can come from the onset of accumulation, the selection of the relevant stimulus or the step from commitment to the press. At a given age, longer non-decision times went with a later CPP half-peak but not with a longer interval from peak to press. A later half-peak fits a later onset of accumulation as well as a slower early rise, and target selection signals modulate both properties in adults (Loughnane et al., 2016). Part of what matures in early adolescence is thus how soon the accumulation signal rises once a target appears. The latency of the P300, which shares its properties with the CPP (Twomey et al., 2015), likewise shortens from childhood into adolescence (van Dinteren et al., 2014).

These schedules join two earlier modelling studies into one trajectory. In 6-to 12-year-olds, the best model required age effects on drift rate and boundary separation but not on non-decision time (Manning et al., 2021), whereas in early adolescence non-decision time was among the components that changed most (Frogner et al., 2026). Each study covered only part of the age range; across 5 to 21 years in one task, the trajectory was staged: drift rate changed early, whereas non-decision time changed little before 10 years and shortened mainly between 11 and 14 years. The results also qualify the view that a single global speed factor drives cognitive development (Kail, 1991; Kail & Ferrer, 2007): when slow accumulation ends in a miss, the early gain appears in accuracy, and speed follows years later. Performance that looks unitary can be assembled from components with distinct trajectories (Coughlin et al., 2024; Ferrand et al., 2020), and speed matures late in other domains too: children select which responses to stop by early preadolescence, but the speed of stopping keeps improving into young adulthood (Albert et al., 2022). Across adulthood, too, speed and the rate of accumulation follow different age courses: in 1.2 million participants, responses slowed from about 20 years through greater caution and slower non-decision processes, whereas drift rate declined only after about 60 years (von Krause et al., 2022).

On a contrast ramp, the timing of the CPP carries more developmental information than its slope. A child with a higher drift rate reaches the bound earlier on the ramp, while the momentary evidence is still weak, so the slope just before the response is set more by the bound than by drift rate, unlike at fixed motion coherence (Manning et al., 2021).

We examined explanations that do not involve the decision components. The SSVEP evidence was sensitive enough to register a developmental change: it followed the physical contrast, was reliable at every age and related to lapses among children of the same age. The lag between drift rate and non-decision time did not arise from trade-offs between parameters, because it persisted with boundary separation fixed, simulated children whose non-decision time matured with their drift rate yielded almost no lag, and the fastest responses changed years after accuracy without any model. Nor did practice, fatigue or psychopathology produce the age effects, which were present from the first run of the task (Supplementary Table S1) and unchanged when psychopathology was controlled.

The study has three limitations. First, the data are cross-sectional, so the schedules capture differences between age groups rather than change within a child; whether an individual child’s non-decision time shortens mainly in early adolescence requires longitudinal data. Second, the participants were recruited through community referral, often because of concerns about psychiatric symptoms (Alexander et al., 2017), and the public data do not include diagnoses or medication. Third, few participants were 18 years or older, so the curves describe development up to late adolescence.

These results narrow where developmental research on perceptual decisions needs to look: at separate changes in how readily children take up registered evidence in middle childhood and how soon the accumulation signal rises in early adolescence, rather than at a single age-related speeding or at early visual encoding. Longitudinal measurements will show whether drift rate and lapse rate mature together within a child and whether non-decision time changes years later. Separating target selection, accumulation onset and motor preparation, each of which has its own neural signature in this family of tasks (Loughnane et al., 2016; O’Connell et al., 2012), will identify the stage that carries the adolescent shortening of non-decision time. The same components matter clinically: children with attention-deficit/hyperactivity disorder differ from their peers mainly in drift rate (Karalunas et al., 2014) and in neural signals of attentional selection and evidence accumulation (Biabani et al., 2025), and in dyslexia, perceptual decision deficits occur largely independently of phonological ones (O’Brien & Yeatman, 2021). Schedules for each component give such clinical comparisons a developmental reference.

## Conclusion

Between early childhood and late adolescence, children became accurate years before they became fast, and neither change was carried by a stronger early visual response. Accuracy rested on fewer lapses and more efficient accumulation of evidence, which matured together in middle childhood. The fastest responses rested on a shorter non-decision time, which matured in early adolescence and went with the timing of the accumulation signal. Perceptual decisions thus develop through components that follow their own schedules after early visual encoding. Each schedule is a developmental reference for its component.

## Data availability statement

The data are the Healthy Brain Network EEG releases 1 to 10 (HBN-EEG), openly available on OpenNeuro under a CC BY-SA 4.0 licence (https://doi.org/10.18112/openneuro.ds005505.v1.0.1, https://doi.org/10.18112/openneuro.ds005506.v1.0.1, https://doi.org/10.18112/openneuro.ds005507.v1.0.1, https://doi.org/10.18112/openneuro.ds005508.v1.0.1, https://doi.org/10.18112/openneuro.ds005509.v1.0.1, https://doi.org/10.18112/openneuro.ds005510.v1.0.1, https://doi.org/10.18112/openneuro.ds005511.v1.0.1, https://doi.org/10.18112/openneuro.ds005512.v1.0.1, https://doi.org/10.18112/openneuro.ds005514.v1.0.1, https://doi.org/10.18112/openneuro.ds005515.v1.0.1). The code for all analyses and figures will be made publicly available upon journal publication.

## Ethics statement

The Healthy Brain Network study was approved by the Chesapeake Institutional Review Board, with written informed consent from adult participants or legal guardians and written assent from minors. The present study analysed only these publicly available, de-identified data and collected no new data, so no further ethical approval was required.

## Permission to reproduce material from other sources

No material is reproduced from other sources.

## Acknowledgements

We thank the participants and families of the Healthy Brain Network and the Child Mind Institute for making the data openly available. Claude (Anthropic) was used to assist with checking the analysis code.

