## supplementary_information for "Children become accurate years before they become fast: electroencephalography dates the separate maturation of decision components"

### ***Supplementary Methods***

#### *Task*

The task was a two-alternative version of the gradual contrast change paradigm (Langer et al., 2017; O'Connell et al., 2012). Participants fixated the centre of an annulus (inner radius 1°, outer radius 6°) formed by two overlaid gratings tilted 45° to the left and to the right of vertical, which phase-reversed at 20 and 25 Hz, respectively. Between targets, both gratings had a contrast of 50%. On each target, the contrast of one grating rose linearly to 100% over 1.6 s while the other fell to 0%, and both returned to 50% over the next 0.8 s. Participants pressed the left button with the left hand when the left-tilted grating increased in contrast and the right button with the right hand when the right-tilted grating did. Each run of 3.1 min contained 12 targets of each type in random order, separated by 2.8, 4.4 or 6.0 s, and a smiley or sad face gave feedback for 400 ms after each target (Langer et al., 2017). Most participants completed three runs, and the median child saw 72 targets.

The response to a target was the first button press between its onset and the next target that the task scored as correct or incorrect; targets without such a press were counted as missed. Accuracy was the proportion of correct responses among all targets, so that missed targets lowered accuracy, and RT analyses used correct responses between 0.2 and 2.5 s.

#### *EEG recording and preprocessing*

EEG was recorded at 500 Hz with a 128-channel HydroCel Geodesic Sensor Net (Electrical Geodesics Inc., Eugene, OR, USA) referenced to Cz (Langer et al., 2017). Each run was processed separately. Channels with a robust z score of log variance above 3, computed from the median and the median absolute deviation across channels, were interpolated with spherical splines. The data were then re-referenced to the average and transformed to current source density (Perrin et al., 1989), which sharpens the spatial separation of centro-parietal and occipital signals. For the CPP, the data were band-pass filtered between 0.1 and 30 Hz and resampled to 100 Hz; for the SSVEP, they were high-pass filtered at 0.1 Hz with a 60-Hz notch filter.

Eye movements and blinks were detected from periocular channels, band-pass filtered between 0.5 and 20 Hz. The vertical derivation was the mean of E25 − E127 and E8 − E126, and the horizontal derivation was E125 − E128. A trial contained an eye event when either peak-to-peak amplitude between 0 and 0.9 s after target onset exceeded the child's median by more than three robust standard deviations, and a trial was flagged as an amplitude artefact when its largest absolute CPP value exceeded the child's median by more than five robust standard deviations. All comparisons between trials of the same child used only trials without eye events.

### *SSVEP evidence*

We averaged five occipital channels (E70, E74, E75, E82 and E83) and computed short-time Fourier transforms with a 400-ms Hann window advanced in 50-ms steps, which gives a frequency resolution of 2.5 Hz with bins centred on 20 and 25 Hz. On each trial, SSVEP evidence was the log ratio of the amplitude at the frequency of the grating that increased in contrast to the amplitude at the frequency of the grating that decreased, minus its mean before target onset. Because it is a ratio, this measure indexes the differential sensory evidence available for the choice independently of the overall size of the visual response (Norcia et al., 2015). Each child's SSVEP evidence was the mean over valid trials between 0.8 and 1.2 s after target onset. Split-half reliability was the Pearson correlation between odd and even trials, corrected with the Spearman–Brown formula, and the onset of the SSVEP evidence was the time at which the average of an age band first reached 25% of its maximum.

### *Centro-parietal positivity*

The CPP was the average of five centro-parietal channels (E54, E61, E62, E78 and E79). Target-locked epochs extended from −0.2 to 2.5 s around target onset and response-locked epochs from −1.0 to 0.3 s around the response, both corrected with the mean between −0.2 s and target onset. To compare children whose recordings differed in noise, we expressed the CPP in units of each child's trial-to-trial noise, the robust standard deviation across trials of the response-locked CPP in the 100 ms before the response. The response-locked CPP slope was the slope of a line fitted to the child's average response-locked CPP between −0.5 and −0.1 s. On single trials, the build-up of the CPP was the slope of the target-locked CPP from 0.2 s after target onset to 0.1 s before the median RT of the child's fastest third of correct trials.

The timing of the CPP was the time at which the target-locked average, smoothed with a 100-ms moving average, first reached half of its maximum between 0.2 and 1.8 s after target onset, with linear interpolation between samples. Single-child averages are too noisy for this criterion, so we retrieved individual latencies by jackknifing within age bands (Miller et al., 1998; Smulders, 2010). Each child's latency was $nJ - (n - 1)J_{-i}$, where n is the number of children in the band, J the half-peak time of the band average and $J_{-i}$ that of the band average without the child. As a second latency measure, we used the time at which half of the positive area between 0.2 and 1.8 s was reached (Kiesel et al., 2008; Liesefeld, 2018).

### *Decision model*

We fitted each child's behaviour with a diffusion model adapted to the gradual targets (Ratcliff & McKoon, 2008). The momentary drift was a drift rate v multiplied by the physical contrast difference, which rose linearly from 0 to 1 over 1.6 s and returned to 0 by 2.4 s. Evidence accumulated with unit noise between symmetric bounds at ±a, and reaching the bound of the grating that increased in contrast produced a correct response. A response was made only if the decision time plus a non-decision time Ter was shorter than 2.5 s. A lapse rate λ captured targets on which no decision was made, whatever the evidence (Ratcliff & Tuerlinckx, 2002). The model therefore predicted correct responses with probability $(1 - \lambda)Pc$, errors with probability $(1 - \lambda)Pe$ and missed targets with probability $\lambda + (1 - \lambda)Pm$, where Pc, Pe and Pm are the probabilities of the diffusion process, together with the quantiles of correct RT.

Parameters were estimated by grid search on a table of simulated predictions, with 4,000 trials per cell and a time step of 10 ms. The grid covered v from 0.3 to 20 in 40 logarithmic steps, a from 0.3 to 4.5 in 43 steps, Ter from 0.10 to 1.60 s in 20-ms steps and λ from 0 to 0.40 in steps of 0.02. The fit minimized the squared error of the 10th, 30th, 50th, 70th and 90th percentiles of correct RT plus four times the squared error of the three response probabilities. Error RTs, which most children produced on few trials, were not fitted. We refitted the data with Ter fixed at 0.5 s and, separately, with a fixed at the sample median, to test whether the age effects on v and Ter depended on trade-offs between parameters. Parameter recovery, predictions of the fitted model and the number of estimates at the grid limits are reported in Supplementary Figures S1 and S2 and Supplementary Table S2.

To test whether the fitting procedure creates a lag between the developmental schedules of drift rate and non-decision time, we simulated every child with its own number of targets and its fitted v, a and λ. Non-decision time followed either the observed age curve or an early curve with the same change from 6 to 16 years and the timing of the drift rate. The simulated data were refitted with the same procedure and the half-change ages recomputed, with five independent random seeds and 500 bootstrap resamples. In a third scenario, non-decision time did not change with age. The contributions of each parameter to the age differences in behaviour were estimated with counterfactual predictions and a Shapley decomposition (Shorrocks, 2013), as described in Supplementary Note S1.

#### *Developmental curves*

Age effects were Spearman correlations with age. To describe when each measure changed, we fitted a natural cubic spline of age with 4 degrees of freedom and knots at the 20th, 40th, 60th and 80th percentiles of age to children aged 5 to 17 years, the range in which the sample was dense, and obtained 95% intervals from 2,000 bootstrap resamples of children. From each curve we derived the half-change age, the age at which the curve had covered half of its change between 6 and 16 years, and the yearly rate of change on a 0.25-year grid, together with the ages at which its interval excluded zero. Differences between half-change ages were computed within each bootstrap sample. The fastest responses were indexed by the 10th percentile of each child's correct RT. Curves with other degrees of freedom and anchor ages, and penalized splines, are reported in Supplementary Table S4.

#### *Relations between measures*

Relations among children of the same age were partial Spearman correlations. Both variables were rank-transformed and residualized on a natural spline of log age with 4 degrees of freedom, sex, release, the number of valid EEG trials, the CPP noise level and the number of interpolated channels, and the residuals were correlated; analyses of CPP timing also controlled age band and log median RT. Intervals came from 2,000 bootstrap resamples of children.

For the single-trial analysis, we took each child with at least 30 correct trials without eye events and regressed log RT on CPP build-up and SSVEP evidence on the same trials, with all three variables z-scored within the child. The regression weights were averaged across children and tested with one-sided Wilcoxon signed-rank tests, and the interval for the difference between the two weights came from bootstrap resampling of children. The relation between coupling and age was the Spearman correlation between each child's CPP weight and age.

For missed targets, we took children with at least three missed and 15 correct trials without eye events. SSVEP evidence between 0.8 and 1.2 s and the mean target-locked CPP between 0.5 and 1.5 s were averaged separately over correct trials, missed targets and the correct trials in the child's lowest third of SSVEP evidence. The ratio of SSVEP evidence on missed to correct trials received a bootstrap interval across children, and the CPP on weak-signal correct trials was compared with the CPP on missed targets with a one-sided Wilcoxon signed-rank test.

In a second analysis, weak-signal trials were selected and measured in separate windows. Each child's lowest third of correct trials was selected by SSVEP evidence in frames centred between 0.6 and 0.8 s, and SSVEP evidence was measured in frames centred between 1.2 and 1.4 s; the reverse direction swapped the two windows. With 400-ms frames, the two windows do not overlap. SSVEP evidence was corrected with the child's mean pre-target log ratio for each target side rather than with each trial's own baseline, so that no trial-level quantity entered both windows. Trials with eye events between 0 and 0.9 s or between 0.9 and 1.6 s were excluded, which left 871 children with at least three missed and 15 correct trials. Differences between weak-signal correct trials and missed targets received 95% bootstrap intervals across children (Supplementary Table S7).

To time the step from commitment to the press, we located the peak of each age band's average response-locked CPP on correct trials without eye events, smoothed with a 100-ms moving average, between 0.5 s before and 0.25 s after the response. Individual peak-to-press intervals were jackknifed within age bands as for the half-peak time and related to non-decision time with the same covariates. Their split-half reliability between odd and even trials was 0.55. Differences between age bands received intervals from 500 bootstrap resamples of children.

### *Scalp maps*

For the scalp maps (Supplementary Figure S3), every channel was processed as for the CPP (0.1–30 Hz, current source density) and as for the SSVEP evidence (0.1-Hz high-pass filter, 60-Hz notch filter, current source density, 400-ms frames). SSVEP evidence was the mean between 0.8 and 1.2 s minus the mean between −0.5 and 0 s, and the CPP was averaged in 0.2-s windows in units of the child's trial-to-trial noise. Maps were averaged over children within each age band who had at least 20 correct trials without eye events, and over the children with at least three missed and 15 correct trials for the comparison of missed and correct targets. The 30 electrodes on the face and neck were left out, and the maps were interpolated with thin-plate splines inside the head outline.

### *Statistical tests*

The main text reports the pooled sample, and Supplementary Table S6 reports releases 1 to 4 and 5 to 10 separately, analysed with identical code. The main tests (Table 2) used one-sided $\alpha = 0.05$ and bootstrap intervals from 5,000 resamples of children. For the prediction that SSVEP evidence does not increase with age, we set an equivalence bound of $\rho = 0.10$ (Lakens, 2017), and the prediction held if the upper limit of the 95% interval of the age correlation lay below this bound. Test 5 compared the fastest and slowest thirds of each child's correct trials without eye events, using the within-child Cohen's d of CPP build-up and of SSVEP evidence. With 1,505 children, a one-sided correlation test at $\alpha = 0.05$ had 80% power to detect $\rho = 0.06$.

### *Psychopathology and sex*

The four psychopathology scores of HBN-EEG, a general factor and specific attention, internalizing and externalizing factors, come from a bifactor model of questionnaire data (Shirazi et al., 2024). We related each score to the decision components with partial Spearman correlations, using the covariates listed above, and controlled the false discovery rate across scores and components (Benjamini & Hochberg, 1995). Associations found in both sets were also estimated in the two sets combined. We repeated the age analyses with attention problems and the general factor as additional covariates, and in children who scored below the median on both. Developmental timing in girls and boys was compared by fitting the curves separately to each sex in the two sets combined.

Analyses were run in Python 3.11 with MNE-Python 1.13.2, NumPy 2.4.6, SciPy 1.17.1, pandas 2.2.3 and statsmodels 0.15.0.

### *Supplementary Note S1. How the decision components share the behavioural change*

We simulated each outcome with the parameters of 5- to 7-year-olds and replaced one parameter at a time by its value in 14- to 17-year-olds, averaging over all orders of replacement (Shapley decomposition; 20,000 common-random-number trials). We did this with the group medians and with the group means of the fitted parameters. At the level of individual children, we predicted each child's median correct RT with one parameter set to the child's own value and the others at the sample median, both for the main fit and for a refit with the decision boundary fixed across children.

| Outcome | Version | Drift rate | Boundary | Non-decision time | Lapse rate |
|---|---|---|---|---|---|
| Accuracy | group medians | 24% | 7% | 7% | 62% |
| Accuracy | group means | 52% | 0% | 11% | 37% |
| Missed targets | group medians | 11% | −3% | 9% | 84% |
| Missed targets | group means | 47% | −11% | 14% | 50% |
| Median correct RT | group medians | 34% | −21% | 87% | – |
| Median correct RT | group means | 67% | −33% | 67% | – |
| RT age slope, children | free boundary | 36% | – | 100% | – |
| RT age slope, children | boundary fixed | 46% | – | 53% | – |

Shares of the change from 5–7 to 14–17 years. Negative shares mean the parameter change worked against the observed change; shares need not sum to 100% for the child-level rows. The two components that carried accuracy, lapse rate and drift rate, did so in every version, whereas the split of speeding between drift rate and non-decision time depended on how the boundary was treated.

### *Supplementary Table S1. Age effects in the first run and in later runs*

| Measure | ρ with age, run 1 | ρ with age, later runs | ρ(age, change from run 1 to later runs) |
|---|---|---|---|
| Accuracy | 0.49 | 0.54 | −0.009 |
| Missed targets | −0.55 | −0.57 | 0.012 |
| Median correct RT | −0.33 | −0.39 | – |

n = 1,506. Mean accuracy rose from 75% in run 1 to 79% in later runs, equally at all ages.

### *Supplementary Table S2. Estimates at a limit of the parameter grid*

| n | At a grid limit | ρ with age, all (v / a / Ter / λ) | ρ with age, without limit estimates |
|---|---|---|---|
| 1534 | 127 (drift 72, lapse 24, Ter 46, boundary 2) | 0.33 / 0.17 / −0.26 / −0.47 | 0.28 / 0.19 / −0.32 / −0.48 |

Grid limits: drift rate 0.3 (lowest value), lapse rate 0.40 (highest), non-decision time 0.10 or 1.60 s, boundary 4.5. Children at the drift floor mostly responded near chance; children at the lapse ceiling missed about half of the targets.

### *Supplementary Table S3. Same-age relation between CPP timing and non-decision time*

| Latency measure | n | Estimates outside 0–2.5 s | Partial ρ with Ter: covariates | + age band | + age band + log RT |
|---|---|---|---|---|---|
| Half-peak time | 1504 | 3.1% | 0.18 | 0.19 | 0.13 |
| Fractional-area latency | 1504 | 1.0% | 0.16 | 0.16 | 0.10 |

Individual latencies were retrieved by jackknifing within age bands (Smulders, 2010) on eye-movement-free correct trials. Covariates as in Table 2. Spearman-based partial correlations are insensitive to the few extreme retrieved values. The same latencies were unrelated to drift rate at a given age.

The interval from the peak of the band-average response-locked CPP to the press, jackknifed in the same way (split-half reliability 0.55), was unrelated to non-decision time at a given age (partial ρ with age band and log RT = −0.05, 95% CI [−0.10, 0.01], n = 1,431). Between 5–7 and 14–21 years, this interval shortened by 100 ms [20, 140] and non-decision time by 224 ms [176, 275].

### *Supplementary Table S4. Lag of non-decision time behind drift rate under alternative curve settings*

| Curve setting | Lag in half-change age, years [95% CI] |
|---|---|
| Natural spline, 3 df | 3.48 [2.23, 4.56] |
| Natural spline, 4 df (main) | 3.69 [2.57, 4.69] |
| Natural spline, 5 df | 3.84 [2.39, 5.08] |
| Natural spline, 6 df | 3.83 [2.31, 5.23] |
| Anchors 7 and 15 years | 3.24 [2.13, 3.96] |
| Anchors 6 and 17 years | 3.54 [2.16, 4.91] |
| Penalized spline (GCV) | 3.59 [2.32, 4.73] |
| Without grid-limit estimates | 3.42 [2.13, 4.32] |

Half-change age: the age at which the fitted curve has covered half of its change between the two anchor ages (6 and 16 years unless stated). 2,000 bootstrap resamples of children. With the boundary fixed across children (500 resamples) the lag was 4.03 years [2.73, 5.33]. Simulated children whose non-decision time did not change with age gave recovered lags of −1.2 to 2.6 years. In behaviour, accuracy among answered targets completed half of its change 4.73 years [3.55, 6.11] before the fastest responses, and accuracy 3.22 years [2.32, 3.97] before median correct RT.

### *Supplementary Table S5. Psychopathology scores and decision components at a given age*

| Score | Measure | Partial ρ (p) |
|---|---|---|
| Attention problems | Lapse rate | 0.061 (0.018) |
| Attention problems | Missed targets | 0.063 (0.016) |
| Attention problems | Drift rate | −0.017 (0.510) |
| Attention problems | Non-decision time | 0.010 (0.690) |

| | | |
|---|---|---|
| Attention problems | Boundary | −0.023 (0.380) |
| Attention problems | SSVEP evidence | −0.027 (0.290) |
| Attention problems | CPP slope | −0.055 (0.034) |
| General psychopathology | Lapse rate | 0.026 (0.330) |
| General psychopathology | Missed targets | 0.041 (0.110) |
| General psychopathology | Drift rate | −0.045 (0.082) |
| General psychopathology | Non-decision time | −0.010 (0.710) |
| General psychopathology | Boundary | 0.007 (0.780) |
| General psychopathology | SSVEP evidence | −0.025 (0.340) |
| General psychopathology | CPP slope | −0.010 (0.710) |
| Internalizing | Lapse rate | −0.059 (0.022) |
| Internalizing | Missed targets | −0.073 (0.005) |
| Internalizing | Drift rate | 0.017 (0.510) |
| Internalizing | Non-decision time | 0.014 (0.590) |
| Internalizing | Boundary | −0.009 (0.730) |
| Internalizing | SSVEP evidence | −0.013 (0.630) |
| Internalizing | CPP slope | 0.011 (0.670) |
| Externalizing | Lapse rate | 0.042 (0.110) |
| Externalizing | Missed targets | 0.041 (0.120) |
| Externalizing | Drift rate | −0.031 (0.230) |
| Externalizing | Non-decision time | −0.007 (0.790) |
| Externalizing | Boundary | 0.026 (0.310) |
| Externalizing | SSVEP evidence | −0.045 (0.083) |
| Externalizing | CPP slope | −0.023 (0.380) |

Partial Spearman correlations with the covariates of Table 2, two−sided p values. Attention problems related to the lapse rate at 0.061 [0.010, 0.112]. Age effects with attention problems and the general psychopathology factor controlled: drift rate 0.32, non−decision time −0.25, lapse rate −0.46. In children below the median on both scores: drift rate +0.337, lapse rate −0.507, non−decision time −0.316. Girls minus boys in half−change age: non−decision time +0.3 years [−1.6, 2.5], drift rate −0.9 years [−2.2, 0.3].

### *Supplementary Table S6. Main tests in releases 1–4 and 5–10*

| Test | Prediction | Releases 1–4 (n = 556) | Releases 5–10 (n = 978) |
|---|---|---|---|
| 1 | Drift rate rises with age | ρ = 0.36; fixed Ter 0.42 | ρ = 0.31; fixed Ter 0.34 |
| 2 | SSVEP evidence does not rise with age and dissociates from drift rate | ρ = −0.13 [−0.21, −0.04]; difference [0.36, 0.58] | ρ = −0.07 [−0.13, −0.01]; difference [0.29, 0.47] |
| 3 | Missed targets decrease with age | ρ = −0.63 | ρ = −0.60 |
| 4 | Response-locked CPP slope relates to drift rate at a given age | partial ρ = 0.11, p = 0.006 | partial ρ = 0.08, p = 0.009 |
| 5 | Fast and slow trials differ more in CPP than in SSVEP evidence | d = 0.22 vs 0.00, p < 0.001 | d = 0.16 vs 0.08, p < 0.001 |
| 6 | Missed targets keep their visual response | 0.76 [0.66, 0.86] | 0.77 [0.69, 0.85] |
| 7 | Missed targets lack the accumulation signal | p < 0.001 | p = 0.006 |

| | | | |
|---|---|---|---|
| 8 | Fastest RT matures after accuracy | 4.60 years [2.50, 6.43] | 4.55 years [3.12, 6.27] |
| 9 | Non-decision time matures after drift rate | 3.36 years [0.34, 4.86] | 3.75 years [2.01, 5.25] |
| 10 | Later CPP half-peak goes with longer non-decision time | partial ρ = 0.13 | partial ρ = 0.14 |

Same tests and covariates as Table 2; rows 8 and 9 give lags in half-change age with 95% bootstrap intervals, row 10 the partial correlation controlling age band and log RT. The main tests were set up on releases 1–4 before releases 5–10 were analysed.

#### *Supplementary Table S7. Weak-signal correct trials selected and measured in separate windows*

| Measure | Selected 0.6–0.8 s, measured 1.2–1.4 s | Selected 1.2–1.4 s, measured 0.6–0.8 s |
|---|---|---|
| SSVEP evidence, missed / correct | 0.80 [0.75, 0.86] | 0.76 [0.66, 0.86] |
| SSVEP evidence, weak / correct | 0.70 [0.64, 0.75] | 0.43 [0.31, 0.54] |
| SSVEP evidence, weak − missed | −0.05 [−0.09, −0.01] | −0.08 [−0.12, −0.04] |
| CPP, missed / correct | 0.15 [−0.26, 0.56] | 0.15 [−0.27, 0.56] |
| CPP, weak / correct | 0.89 [0.63, 1.13] | 0.74 [0.44, 0.99] |
| CPP, weak − missed | 0.06 [0.02, 0.11] | 0.05 [0.00, 0.10] |
| Response-locked CPP, weak / correct | 0.97 [0.87, 1.07] | 0.95 [0.83, 1.05] |
| Median RT, weak-signal trials (s) | 1.70 | 1.70 |

n = 871 children with at least 3 missed and 15 correct trials free of eye events. Weak-signal trials are each child's lowest third of correct trials by SSVEP evidence in the selection window; SSVEP evidence is reported in the other window. The CPP is the mean between 0.5 and 1.5 s after target onset (response-locked, between 0.1 s before and the response) in units of the child's trial-to-trial noise. Ratios are means across children; brackets give 95% bootstrap intervals. Within children, SSVEP evidence in the two windows correlated weakly across correct trials (median ρ = 0.11); median RT on all correct trials was 1.69 s. Three quarters of these children were younger than 11 years, the ages at which missed targets were most frequent.

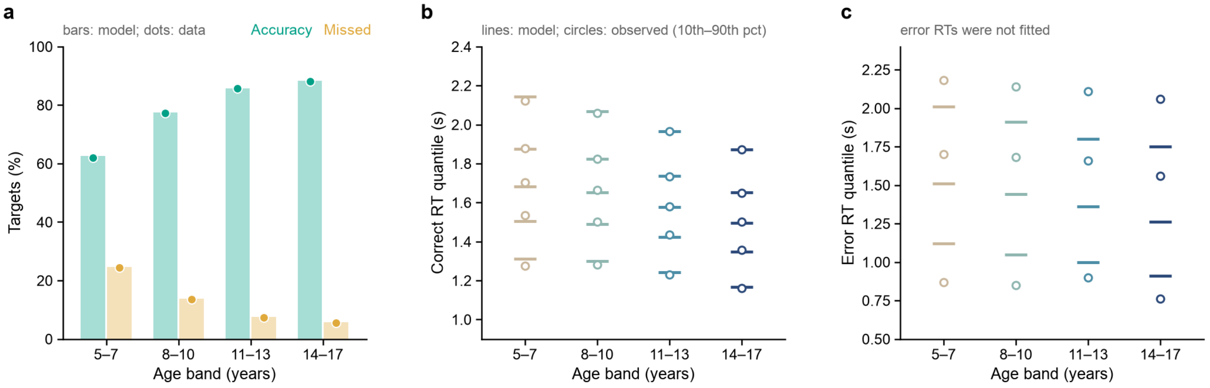


**Supplementary Figure S1. Model predictions against the data by age band.** (a) Accuracy and missed targets predicted by the fitted model (bars) and observed (dots). (b) Quantiles of correct RT (10th, 30th, 50th, 70th, 90th percentiles): model (lines) and data (circles). (c) Quantiles of error RT (10th, 50th, 90th percentiles), which the model was not fitted to; observed errors include more fast responses and a slower median than predicted. Predictions use each child's fitted parameters.

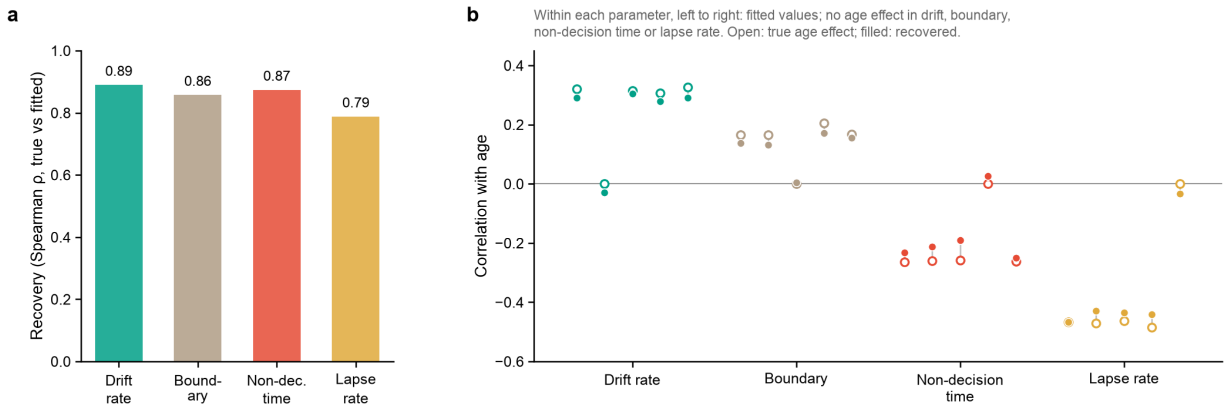


**Supplementary Figure S2. Recovery of parameters and of their age effects.** Children were simulated with their own trial counts and fitted parameters and refitted with the same model. (a) Spearman correlation between true and recovered parameters. (b) Age effects of each parameter, true (open) and recovered (filled), when the simulated parameters kept their fitted values and when the age effect of one parameter at a time was removed. Removing an age effect brought its recovered value close to zero, so age effects did not transfer from one parameter to another.

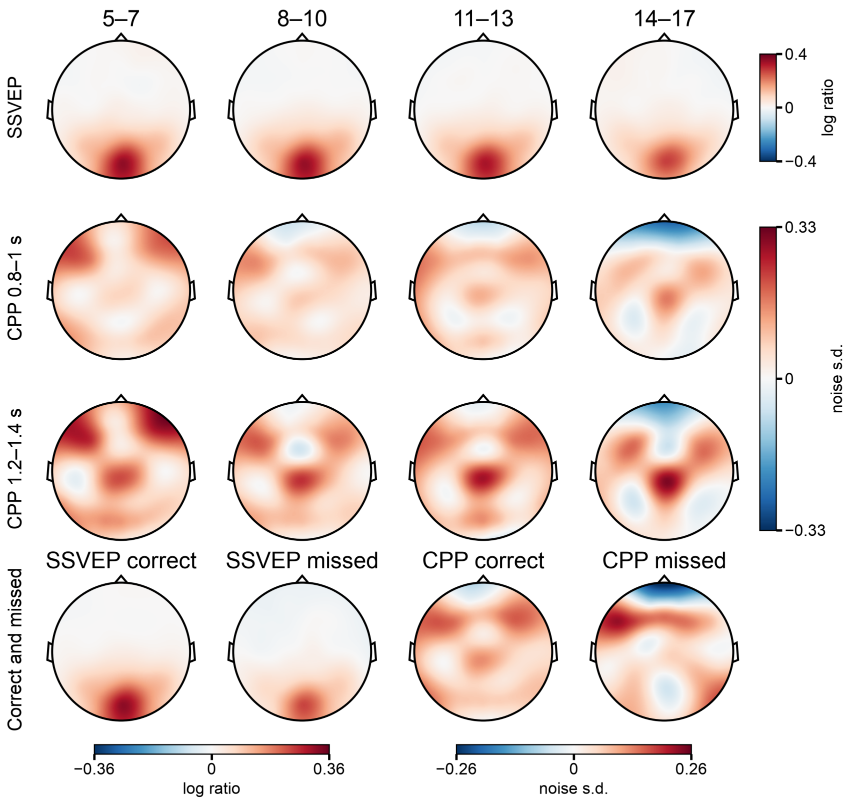


**Supplementary Figure S3. Scalp maps of the visual response and the accumulation signal.** Group means of SSVEP evidence (0.8–1.2 s after target onset) and of the target-locked CPP in two windows on correct trials, by age band (top three rows; 294, 552, 338 and 210 children), and of both signals on correct trials and missed targets (0.5–1.5 s for the CPP; bottom row; 956 children). Red, positive; blue, negative. The SSVEP row has its own colour scale and the CPP rows share one; in the bottom row, each signal has its own scale. Measured against neighbouring frequencies before the target, the SSVEP itself rose slightly with age ($\rho = 0.06$, $p = 0.03$), whereas its raw amplitude fell ($\rho = -0.20$). SSVEP, steady-state visual evoked potential; CPP, centro-parietal positivity.

### *Supplementary references*


Benjamini, Y., & Hochberg, Y. (1995). Controlling the False Discovery Rate: A Practical and Powerful Approach to Multiple Testing. Journal of the Royal Statistical Society Series B: Statistical Methodology, 57(1), 289–300. https://doi.org/10.1111/j.2517-6161.1995.tb02031.x

Kiesel, A., Miller, J., Jolicœur, P., & Brisson, B. (2008). Measurement of ERP latency differences: A comparison of single-participant and jackknife-based scoring methods. Psychophysiology, 45(2), 250–274. https://doi.org/10.1111/j.1469-8986.2007.00618.x

Lakens, D. (2017). Equivalence Tests. Social Psychological and Personality Science, 8(4), 355–362. https://doi.org/10.1177/1948550617697177

Langer, N., Ho, E. J., Alexander, L. M., Xu, H. Y., Jozanovic, R. K., Henin, S., Petroni, A., Cohen, S., Marcelle, E. T., Parra, L. C., Milham, M. P., & Kelly, S. P. (2017). A resource for assessing information processing in the developing brain using EEG and eye tracking. Scientific Data, 4(1), 170040. https://doi.org/10.1038/sdata.2017.40

Liesefeld, H. R. (2018). Estimating the Timing of Cognitive Operations With MEG/EEG Latency Measures: A Primer, a Brief Tutorial, and an Implementation of Various Methods. Frontiers in Neuroscience, 12, 765. https://doi.org/10.3389/fnins.2018.00765

Miller, J., Patterson, T., & Ulrich, R. (1998). Jackknife-based method for measuring LRP onset latency differences. Psychophysiology, 35(1), 99–115. https://doi.org/10.1111/1469-8986.3510099

Norcia, A. M., Appelbaum, L. G., Ales, J. M., Cottereau, B. R., & Rossion, B. (2015). The steady-state visual evoked potential in vision research: A review. Journal of Vision, 15(6), 4. https://doi.org/10.1167/15.6.4

O'Connell, R. G., Dockree, P. M., & Kelly, S. P. (2012). A supramodal accumulation-to-bound signal that determines perceptual decisions in humans. Nature Neuroscience, 15(12), 1729–1735. https://doi.org/10.1038/nn.3248

Perrin, F., Pernier, J., Bertrand, O., & Echallier, J. (1989). Spherical splines for scalp potential and current density mapping. Electroencephalography and Clinical Neurophysiology, 72(2), 184–187. https://doi.org/10.1016/0013-4694(89)90180-6

Ratcliff, R., & McKoon, G. (2008). The Diffusion Decision Model: Theory and Data for Two-Choice Decision Tasks. Neural Computation, 20(4), 873–922. https://doi.org/10.1162/neco.2008.12-06-420

Ratcliff, R., & Tuerlinckx, F. (2002). Estimating parameters of the diffusion model: Approaches to dealing with contaminant reaction times and parameter variability. Psychonomic Bulletin & Review, 9(3), 438–481. https://doi.org/10.3758/BF03196302

Shirazi, S. Y., Franco, A., Hoffmann, M. S., Esper, N. B., Truong, D., Delorme, A., Milham, M. P., & Makeig, S. (2024). HBN-EEG: The FAIR implementation of the Healthy Brain Network (HBN) electroencephalography dataset. bioRxiv. https://doi.org/10.1101/2024.10.03.615261

Shorrocks, A. F. (2013). Decomposition procedures for distributional analysis: a unified framework based on the Shapley value. The Journal of Economic Inequality, 11(1), 99–126. https://doi.org/10.1007/s10888-011-9214-z

Smulders, F. T. Y. (2010). Simplifying jackknifing of ERPs and getting more out of it: Retrieving estimates of participants' latencies. Psychophysiology, 47(2), 387–392. https://doi.org/10.1111/j.1469-8986.2009.00934.x
